# Pulse-chase experiments reveal dynamics of RNA binding protein Exuperantia in *Drosophila melanogaster egg* chambers

**DOI:** 10.1101/2022.12.06.519343

**Authors:** Diana V. Vieira, Rita R. Carlota, Jorge de-Carvalho, Ivo A. Telley

## Abstract

In cells, mRNA can be associated with various proteins, forming ribonucleoprotein complexes (RNPs) which take part in spatiotemporal control of translation. In the *Drosophila melanogaster* developing egg chamber, a set of RNPs is transported from the nurse cells to the oocyte and targeted selectively to specific cellular locations. This mRNA sorting process leads to the final oocyte polarization pre-defining the body axes of the future embryo. However, how mRNA is encoded for selection and directed transport is mechanistically not well understood. A master mRNA involved in body axes formation is *bicoid*, which localizes anterolaterally and is essential for head and thorax definition of the embryo. A protein that was identified essential for *bicoid* anterior localization is Exuperantia (Exu). Here, we use a live imaging–based pulse-chase approach, which reveals selective transport dynamics of Exu from nurse cells to the oocyte during mid to late-stage oogenesis.

## Introduction

In vertebrates, the main body axes are defined during early embryonic development (Niehrs, 2010). Conversely, insects define the two main axes prior to fertilization. Body axis determination has been well characterized in *Drosophila melanogaster*, where the localization of *bicoid* (*bcd*) and *oskar* (*osk*) mRNA to the anterior and posterior poles of the oocyte, respectively, defines the primary axis of the embryo (Riechmann and Ephrussi, 2001). RNA localization and translational control target proteins to specific intracellular locations (Blower, 2013). This strategy has the advantage of restricted translation at discrete regions of the cytoplasm, which is central for developmental patterning (Berleth et al., 1988; Martin and Ephrussi, 2009). Local translation of *bcd* and *osk* in the oocyte create protein gradients throughout the later embryo (Gregor et al., 2007). Bicoid protein establishes the anterior pattern of the embryo – head and thorax (Frohnhöfer and Nüsslein-Volhard, 1986) – while Oskar protein retains *nanos* mRNA at the posterior defining abdomen and germline (Lehmann and Nüsslein-Volhard, 1986; Ephrussi et al., 1991).

*bcd* and *osk* mRNAs are synthesized in the nurse cells (NC) and contain localization signals (LEs) that are recognized and consecutively trigger assembly into ribonucleoprotein (RNP) particles prior to nuclear export. These particles undergo further remodeling on their way to the oocyte (Cha et al., 2001). However, the molecular kinetics of this remodeling are unclear. Translocation into the oocyte occurs through ring canals (RC) mediated by an unresolved mechanism involving microtubules (MT), dynein/dynactin and Bic-D/Egl (Kugler and Lasko, 2009). Once RNPs reach the egg cytoplasm, a network of MTs and motor proteins is responsible for the targeted localization of the different mRNAs (St. Johnston, 2005). How one type of mRNA is sorted towards the far-end posterior pole while the other type is enriched at the anterior side proximal to the NCs is an open question. Past studies have put forward a model in which the LEs of these mRNAs are recognized by diverse proteins, transporting them by different MT-based molecular motors and trapping mRNAs at cortical sites (Johnstone and Lasko, 2001; Heredia and Jansen, 2004; Marchand et al., 2012).

Four possible model mechanisms can lead to mRNA targeting: a) local synthesis of the RNA, known to occur in large syncytial cells and in the *Drosophila* oocyte; b) protection from degradation only at places where mRNA is needed (Bergsten and Gavis, 1999); c) diffusion and local removal – the mRNA is enriched in a region while being sequestered by trans-acting elements in another region (Forrest and Gavis, 2003); d) active transport along the cytoskeleton – mRNA travels along actin filaments and MT (Wilhelm and Vale, 1993), using the three families of molecular motors: myosin, dynein, kinesin (St. Johnston, 2005; Clark et al., 2007; Gáspár et al., 2017). The mechanisms are not exclusive as experimental evidence supporting each of these mechanisms is available. Thus, they could be working in concert in the *Drosophila* egg chamber.

Past gene mutagenesis studies indicate that the proteins Exuperantia (Exu), Swallow (Swa) and Staufen (Stau) are important for the anterior localization of *bcd* mRNA (Macdonald et al., 1991; Vogt et al., 2006; Theurkauf and Hazelrigg, 1998). Exu is also present in RNPs containing *osk* mRNA and is required for *osk* posterior localization (Lin et al., 2006). This localization requires Ypsilon-Schachtel (Yps), a cold shock protein of *bcd*. Exu and Yps were shown to co-purify and interact *in vitro* even in the absence of mRNA (Wilhelm et al., 2000). Exu is essential for the proper localization of *bcd* to the anterior (Hazelrigg et al., 1990). This localization revealed to be biphasic: an early phase dependent on Exu phosphorylation by Par-1, and a late phase less dependent on phosphorylation (Riechmann and Ephrussi, 2004). It was first thought to be an exonuclease due to its high sequence homology with other EXO-domain containing proteins. However, the most recent structural and biochemical analysis suggested that Exu could be an RNA-binding protein because it does not contain the characteristic catalytic site of exonucleases (Lazzaretti et al., 2016). Further analysis showed that a large surface area is positively charged, consistent with nucleic acid binding regions, and residues were identified that modulate RNA binding affinity. This structural analysis of Exu fragments also suggested that full-length Exu forms a dimer. Mutagenesis of the putative dimerization domain of endogenous Exu in the fruit fly showed defects in *bcd* localization (Lazzaretti et al., 2016).

Exu’s essential role may be that it is a core component of the assembly pathway and the transport of RNA-protein complexes from NCs to the oocyte. However, the kinetics of Exu binding to RNPs *in vivo* is hitherto unexplored. Specifically, whether its continuous binding to *bcd* is required throughout NC-oocyte transport (co-transport), and if it is encoding RNP sorting or targeting, are still open questions. Here, we report a successful strategy to purify full-length Exu protein constructs (herein called sExu_531_), which we then use in live cell imaging–based pulse– chase experiments to measure Exu movement inside the *Drosophila* egg chamber. Interestingly, our biochemical analysis of sExu_531_ did not indicate dimerization under physiological buffer conditions. Live imaging revealed that during stage 9 and 10A of oogenesis GFP::sExu_531_ is enriched around nurse cell nuclei and then moves rather towards the oocyte than to anterior nurse cells. This suggests that Exu is directionally transported, presumably with associated mRNA.

## Results

### Purification of full-length, fluorescently tagged Exu

An N-terminal GFP fusion construct of Exu was shown to rescue oogenesis of a null mutant allele in *Drosophila* (Wang and Hazelrigg, 1994). Thus, we designed a protein expression construct containing the full coding sequence of the *exu* gene (sExu_531_) and fused it to an N-terminal GFP or mCherry with His_6_ tag sequence for affinity purification (Fig. 1A). Proteins were expressed in *E*.*coli* and purified by following a two-step purification protocol. Cell lysates were first flown over a chelating agarose resin for His-tag mediated affinity and, subsequently, eluted with a linear gradient of imidazole. Next, we performed ionic exchange chromatography to separate the sExu_531_ construct from non-specifically binding biomolecules. The resulting sample was buffer exchanged into a buffer that mimics intracellular physiological conditions (Telley et al., 2013). Finally, we performed size exclusion chromatography to assess potential oligomerization of sExu_531_ constructs, as suggested by earlier literature (Lazzaretti et al., 2016). Some critical steps that helped keeping these Exu protein constructs soluble and stable were as follows: Firstly, the sequence of *exu* required codon optimisation to be expressed in bacterial cell lines. *exu* has 24 forbidden codons in *E. coli*, and expression in *E. coli* Rosetta did not yield a high enough overexpression. After codon optimisation, which was achieved using a synthetic construct (sExu_531_), we found that the *E. coli* strain BL21 yielded the best overexpression of both fusion constructs. Secondly, protein expression was only efficient by IPTG induction at 25ºC for 4 hours; longer incubation periods combined with a range of temperatures and inducer concentrations were tested but led to the formation of inclusion bodies and, consequently, very low yield of Exu proteins in the soluble fraction upon cell lysis. Of note, an auto-induction protocol did not lead to sufficiently high expression. Thirdly, after achieving optimal conditions for protein expression, the cell disruption and stabilization had to be optimized by using a French Press instead of an ultrasound probe. Ultrasound mediated cell disruption in all tested conditions (frequency and pulse time) led to sample degradation. Lastly, optimization of ionic strength and pH during the first purification step proved important, and only cation exchange mediated purification worked successfully. Buffer exchange to physiological (intracellular) proved challenging for His_6_-mCherry::sExu_531_, which presented higher surface adsorbance, but mitigated by overnight soaking of the concentrator membranes (cellulose).

**Figure 1:**
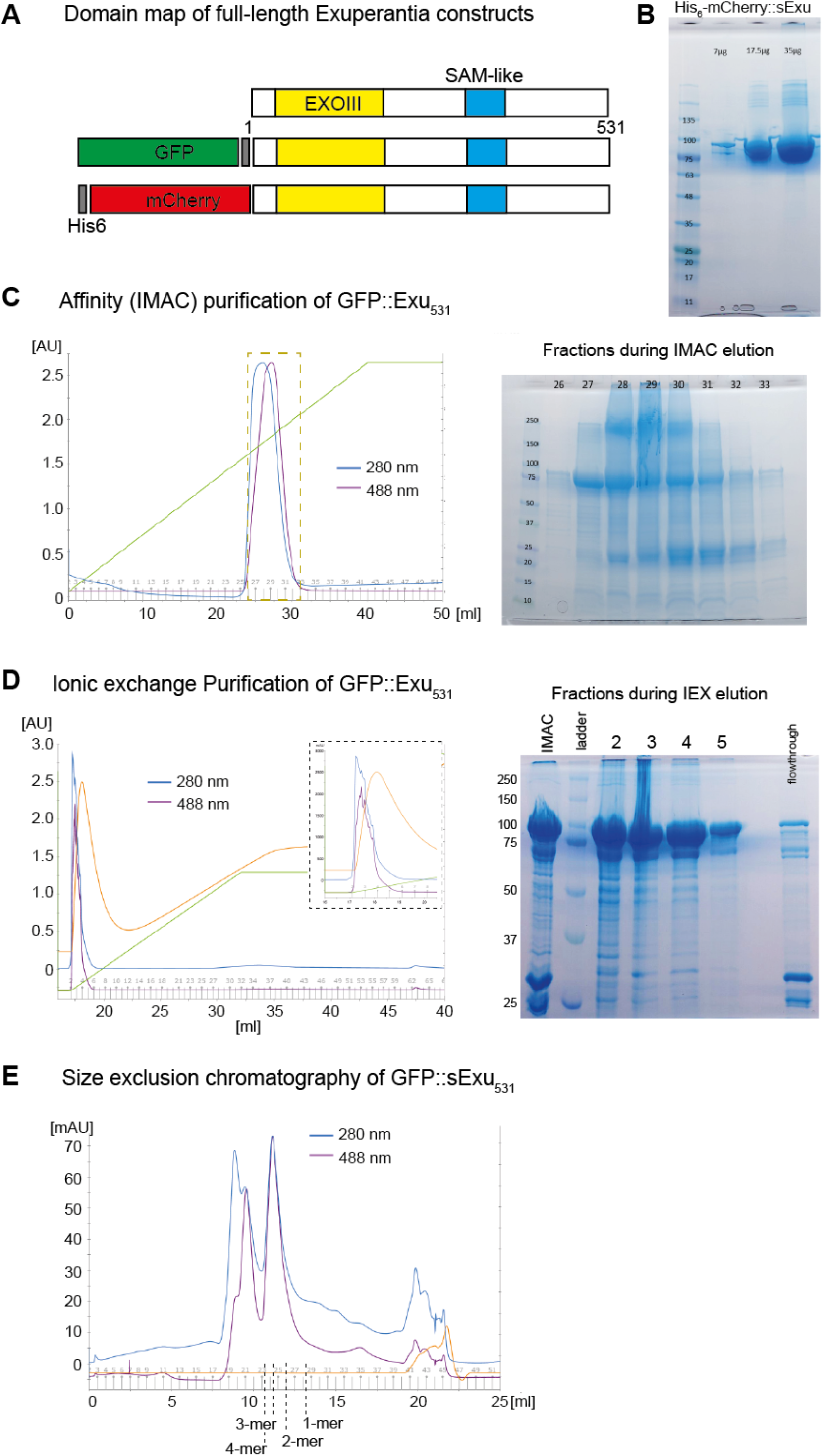
Purification of full-length, fluorescently tagged Exu. **(A)** Sequence domain maps of the native Exu coding sequence (top) with the location of the characteristic EXO-like domain (yellow) and the SAM-like domain (blue). The synthetic constructs with fluorescent fusion protein and hexa-histidine tag are shown below. **(B)** SDS PAGE gel of three loading concentrations of His_6_-mCherry::sExu531 after his-mediated affinity purification, ionic exchange and size exclusion chromatography. The theoretical molecular mass of the construct is 89 kDa. **(C)** A typical Immobilized Metal Affinity Chromatography (IMAC) elution profile using a linear concentration gradient of 0.5 M imidazole (green line). Absorption of 280 nm and 488 nm (GFP) were measured during flowthrough. The dashed rectangle in the profile plot marks the fractions considered for further purification steps, as shown in the SDS PAGE gel on the right. **(D)** Left: Chromatogram during ionic exchange purification (IEX), as described in the Methods section. The green line represents the linear buffer gradient applied during elution. The fractions under the peak of 280 nm and 488 nm absorption (see inset) were analysed on SDS PAGE gel and either used for SEC analysis, frozen for storage or used directly functional experiments. Right: SDS PAGE gel showing samples during a full round of purification, with IMAC elution, fractions 2–5 during IEX, and flowthrough. **(E)** Chromatogram during size exclusion chromatography (SEC) of fraction 4 shown in panel D, using 280 nm and 488 nm absorption. We determined the range of molecular weights using a calibration curve (see Methods). The first peak of the curves suggests existence of larger aggregates, while the second peak represents trimers.

With this approach, and above mentioned optimization steps, we obtained experimentally relevant amounts at concentrations up to 10 mg/ml of purified GFP::sExu_531_ and mCherry::sExu_531_ protein that was stable in buffer at physiological ionic and pH condition (Fig. 1B). Surprisingly, from the three repeats of purifications per construct that we performed, none of the size exclusion chromatography profiles pointed at a robust dimerization of GFP::sExu_531_, at these physiological conditions. The chromatography profiles were compatible with a trimeric or higher oligomerization state (Fig. 1C). However, this was not matching the structural analysis of truncated Exu protein constructs (Lazzaretti et al., 2016). These truncations may favour dimerization and crystal formation, compatible with our so far unsuccessful attempts in obtaining crystals from the full-length protein. Thus, we prefer an interpretation of our and other’s data in which Exu dimerization occurs in a native environment, catalysed by binding of *bcd* mRNA.

### GFP::sExu_531_ moves with posterior bias from nurse cells to the oocyte in stage 9 to 10A

Past studies have suggested that Exu is part of the RNP sorting machinery that transports mRNAs such as *bcd* from nurse cells to the oocyte (Wang and Hazelrigg, 1994; Theurkauf and Hazelrigg, 1998). Whether co-transport of Exu occurs while being permanently bound to the RNP is an open question to address. Addressing this would clarify if Exu is needed for loading of mRNAs to the transport machinery only, or whether it is required throughout the entire transport path. One study reported the temporal change of Exu localisation (Theurkauf and Hazelrigg, 1998). However, analysing signals in constitutively expressing model systems, such as a GFP::Exu transgenic line with abundant fluorescent fusion protein, has the disadvantage that reporter signals cannot be assigned to origin and destination. Because GFP::Exu localises in particles, it provided information about short individual events at best. Thus, we devised an experimental approach to visualize and quantify Exu movement from cell to cell within a living egg chamber. Having the recombinant purified sExu_531_ protein construct tagged with fluorescent reporter at hand, it enabled us to study protein reporter transfer in a ‘pulse-chase’ manner using time-lapse imaging for detection of spatial changes. We injected a range of protein amounts (same volume, different concentrations) of tagged sExu_531_ protein into a single nurse cell by motorised micromanipulation (Fig. 2A,B). For bulk amounts, we then measured the change of fluorescence signal corresponding to this reporter (GFP or mCherry) in neighbouring nurse cells and in the oocyte. While a transgenic fly line expressing full-length Exu tagged with GFP shows particulate localization of GFP::Exu in nurse cells during mid-oogenesis (Fig 2C, top), this localization changed to more homogenous distribution in late stages (Fig 2C, bottom) (Wang and Hazelrigg, 1994). This suggests that Exu is either degraded in nurse cells or transported into the oocyte, and thus diluted overall. Our experiments show that, shortly after injection, GFP::sExu_531_ signal is detectable at the ring canal connecting the injected nurse cell with the oocyte (Fig. 2D,E, Video 1). This marks the entry into the oocyte, where it is diluted hence the signal decreases distal from the ring canal. We also detected GFP signal entering neighbouring nurse cells, but we see a preference for the lateral rather than the anterior nurse cells (Fig. 2E, arrows). This suggests that the cell-to-cell transport is directed or spatially regulated. Notably, we see slow accumulation of GFP::sExu_531_ signal at the anterior end of the oocyte (Fig. 2F) suggesting that the once injected Exu protein is anchored at the anterior end, an established localization site for *bicoid*. This suggests two things: a) our construct is functional and interacts with the transport machinery, b) its destination matches that of *bicoid*. The anterior end–anchoring could be confirmed by injecting GFP::sExu_531_ directly into a stage 10A oocyte, in which cytoplasmic stream occurs. One would expect that streaming would distribute and dilute proteins that remain cytosolic. In contrast, our construct localized at the anterior end after 15–20 min (Fig. 2G), again suggesting that it is anchored there. Overall, we show that the purified GFP::sExu construct moves from nurse cell origin to the oocyte where it is anchored anteriorly.

**Figure 2:**
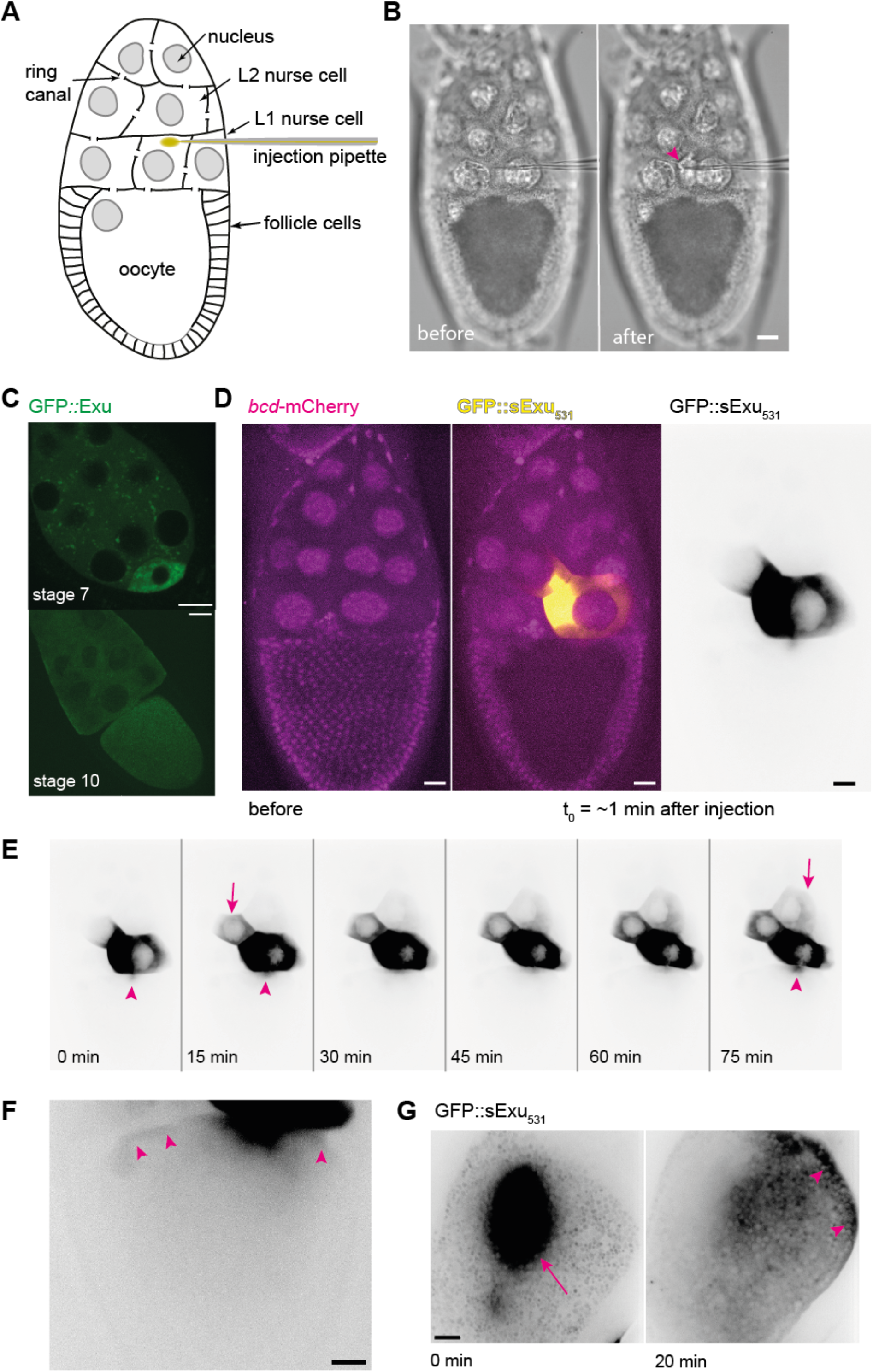
GFP::sExu_531_ dynamics in pulse-chase experiments. **(A)** Schematic showing the experimental design. Living egg chambers from Drosophila are prepared for single cell injection and subsequent time-lapse imaging. A nurse cell that is contacting the oocyte (L1) is selected for injection GFP::sExu_531_ protein. **(B)** Transmission light micrographs of a stage 10A egg chamber before (top) and after (bottom) injection of protein. The change in contrast at the tip of the micropipette (arrowhead) highlights the additional volume inside the nurse cell after injection. **(C)** Maximum intensity z-projection of the fluorescence of egg chambers expressing GFP::Exu constitutively, at different stages. At mid-oogenesis GFP::Exu localizes in particles while in late oogenesis (stage 10) is more homogenous an slightly concentrated at the anterior end of the oocyte. **(D)** Maximum intensity z-projection of the fluorescence of the egg chamber expressing *bcd*MS2(24x) and MCP::mCherry (magenta) before the injection (left), and after injection (midle) overlaid with the maximum intensity z-projection of GFP::sExu_531_ (yellow). The left panel shows the median intensity z-projection, which enhances dimmer signals next to very bright signals, of the GFP channel only with inverted greyscale (black: high intensity, white: low intensity). **(E)** Time lapse of the signal from GFP::sExu_531_ after injection. Within minutes after injection, Exu fills the entire nurse cell volume and travels through the ring canal to the oocyte (arrowhead) where it is diluted and becomes undetectable. Exu is also slowly moving to neighbouring nurse cells located laterally (arrow, t=15min) but much more slowly towards anterior nurse cells (arrow, t=75min). Movement to the oocyte persists throughout the observation. **(F)** A faint crescent of signal is detected at the anterior end of the oocyte, suggesting anchoring of sExu_531_ together with *bicoid* mRNA. **(G)** Injection of GFP::sExu_531_ protein into a stage 10B oocyte that undergoes cytoplasmic streaming (left, arrow). Time-lapse imaging revealed that GFP::sExu531 protein localizes at the anterior end of the oocyte (right, arrowheads), highlighting its recruitment to the anterior, an established localization pattern of *bicoid* mRNA. All scale bars are 20 µm.

## Discussion

This short report provides two new pieces of information about the biochemistry and the biology of Exuperantia protein for mRNA transport in *Drosophila*. Firstly, we show that under conditions mimicking intracellular ionic and pH levels we can produce a stable purified full-length Exu protein and found that this construct forms a trimer or tetramer. We note that the observation of oligomeric state is unlikely influenced by the fluorescent fusion tag, since in our hands a tag-free version of sExu_531_ is showing a similar elution profile during size exclusion chromatography. One possible explanation is that the dimerization of Exu is catalysed by mRNA binding in vivo. Crystal structure analysis of a truncated Exu construct with mutations of the Loop 1 domain suggested that an Exu dimer binds *bicoid* mRNA (Lazzaretti et al., 2016) and it remained unclear if these modifications potentially change the oligomerization state. Further, in vivo analysis may help resolve this conundrum, for example by in vivo reconstitution of Exu– bcd complexes in *Drosophila* ovarian cell extract.

Secondly, and more interestingly, we show that Exu reporter signal moves with a bias along the anterior-posterior axis towards the posterior – the oocyte. After injection, fluorescently tagged Exu initially localizes and concentrates around the nurse cell nucleus and then moves either laterally to the next nurse cell or directly to the oocyte if this cell contact exists. Fluorescence signal movement is not compatible with free or constrained, unbiased diffusion. We conclude that Exu is directionally transported to the oocyte, and that this transport is microtubule dependent because microtubule drugs perturb Exu mobility. (Theurkauf and Hazelrigg, 1998). Consistent with these past studies and adding to them, this data supports the following functional lifetime of Exu in mRNA transport: it becomes part of an RNP close to the nurse cell nucleus, where mRNAs are processed after passing the nuclear envelope and slicing machinery. These particles recruit microtubule-associated molecular motors and move from nurse cell through ring canals to nurse cell or oocyte. Whether the microtubule organization alone generates the anterior-posterior bias is yet to be determined. However, Exu is most certainly not simply a loading factor, but it maintains transport to the destination. Sorting of the anterior and posterior targets may then occur at the anterior end of the oocyte, as it is where all mRNA enters the oocyte by passing through RCs. It is possible that, depending on developmental stage, a member of the complex is modified to engage posterior-directed motor transport with higher probability than anterior anchoring, or vice versa. Recently, molecular sorting by loading and activating kinesin-1 motor was resolved for *oskar* (Gáspár et al., 2017). Whether Exu plays a role in this process remains to be determined and now we have tools in hand to do just that.

## Video Legends

**Video 1:** Median intensity Z-projection from a 3D time-lapse movie of an egg chamber showing the green signal of GFP::sExu_531_ after injection. The black line in the first frame marks the oocyte cortex boundary. Time in min, scale bar 20 µm. Playback frame rate is 10 frames/min. In support of Fig. 2.

## Materials and Methods

### Cloning

#### a) GFP-His_6_::sExu_531_

The plasmid containing the optimized sequence of *Drosophila exu* (1656bp), pHTP9_GFP-His_6_-sExu was obtained from NZYTech, resulting in a final construct of 7663bp corresponding to a recombinant protein containing an N-terminal GFP module followed by an hexa-histidine tag and by a Tobacco Etch virus (TEV) protease recognition sequence, referred to herein as GFP::sExu_531_, with the following sequence:

MGVSKGEELFTGVVPILVELDGDVNGHKFSVSGEGEGDATYGKLTLKFICTTGKLPVPWPTLV TTLTYGVQCFARYPDHMKQHDFFKSAMPEGYVQERTIFFKDDGNYKTRAEVKFEGDTLVNRIE LKGIDFKEDGNILGHKLEYNYNSHKVYITADKQKNGIKVNFKTRHNIEDGSVQLADHYQQNTP IGDGPVLLPDNHYLSTQSALSKDPNEKRDHMVLLEFVTAAGITLGMDELYKSSGPSGSSHHHH HHSSGPQQGLRENLYFQG<u>MVADNIDAGVAIAVADQSSSPVGDKVELPAGNYILVGVDIDTTGR RLMDEIVQLAAYTPTDHFEQYIMPYMNLNPAARQRHQVRVISIGFYRMLKSMQTYKIIKSKSEIAALKDFLNWLEQLKTKAGPSSDGIVLIYHEERKFIPYMILESLKKYGLLERFTASVKSFANSINLAKASIGDANIKNYSLRKLSKILSTTKEEDAACSASTSGSGSGLGSGSSMVSDSVSISPRDSTVTNGDDKQSSKNAVQGKRELFDGNASVRAKLAFDVALQLSNSDGKPEPKSSEALENMFNAIRPFAKLVVSDVLELDIQIENLERQNSFRPVFLNYFKTTLYHRVRAVKFRIVLAENGFDLNTLSAIWAEKNIEGLDIALQSIGRLKSKDKAELLELLDSYFDPKKTTVKPVVKGNSNNNNNYRRRNRRGGRQSVKDARPSSSPSASTEFGAGGDKSRSVSSLPDSTTKTPSPNKPRMHRKRNSRQSLGATPNGLKVAAEISSSGVSELNNSAPPAVTISPVVAQPSPTPVAITASN</u>*

#### b) His_6_-mCherry::sExu_531_

To generate the His_6_-mCherry::sExu_531_ construct we amplified mCherry DNA sequence from pROD17 plasmid (gift from Mariana Pinho’s lab, ITQB) and sExu_531_ DNA sequence from pHTP9_GFP-His_6_::sExu_531_ (described above) and cloned it into pET28a plasmid resulting in a final construct of 7588 bp. Phusion (HF) DNA polymerase was used with a modified PCR reaction (50 µl volume) optimized for this purpose adding the following reagents: 5X Phusion HF buffer, forward primer at 10 µM, reverse primer at 10 µM, 10 mM dNTPs, template DNA at 20 ng/µl, 0.5 µl Phusion HF and nuclease-free water. The following PCR protocol was used: initial denaturation at 98°C for 30 s, then denaturation at 95°C for 10 s, annealing at 72°C for 45 s, extension at 72°C for 45 s for 30 cycles, final extension at 72°C for 5 min and hold at 4°C. The entire reaction volume was used to run a 1% agarose gel electrophoresis. The band corresponding to the size of the sExu_531_ (1656 bp) and mCherry (681 bp) were cut using a scalpel and DNA was purified using e QIAquick PCR Purification Kit according to manufacturer instructions. pET28a plasmid was transformed into chemically competent *E*.*coli* DH5α cells cultured with the appropriate selective conditions (Kanamycin resistance), and a miniprep was performed to extract the DNA (NZYTech kit). The plasmid pET28a and mCherry purified PCR product were enzymatically hydrolysed with NdeI and BamHI-HF restriction enzymes followed, purified from an agarose gel and underwent a T4 ligation reaction. The resulting pET28a-mCherry and sExu_531_ purified PCR product were then enzymatically hydrolysed with EcoRI and XhoI restrictions enzymes, purified from an agarose gel and followed to a T4 ligation reaction, to insert sExu downstream of mCherry. The final construct was sent for sequencing to confirm the quality and presence of the recombinant coding sequence of a fusion protein containing an hexa-histidine tag, followed by the mCherry module upstream of sExu, as follows:

MGSSHHHHHHSSGLVPRGSHMMAIIKEFMRFKVHMEGSVNGHEFEIEGEGEGRPYEGTQTAKL KVTKGGPLPFAWDILSPQFMYGSKAYVKHPADIPDYLKLSFPEGFKWERVMNFEDGGVVTVTQ DSSLQDGEFIYKVKLRGTNFPSDGPVMQKKTMGWEASSERMYPEDGALKGEIKQRLKLKDGGH YDAEVKTTYKAKKPVQLPGAYNVNIKLDITSHNEDYTIVEQYERAEGRHSTGGMDELYKGSEF <u>MVADNIDAGVAIAVADQSSSPVGDKVELPAGNYILVGVDIDTTGRRLMDEIVQLAAYTPTDHFEQYIMPYMNLNPAARQRHQVRVISIGFYRMLKSMQTYKIIKSKSEIAALKDFLNWLEQLKTKAGPSSDGIVLIYHEERKFIPYMILESLKKYGLLERFTASVKSFANSINLAKASIGDANIKNYSLRKLSKILSTTKEEDAACSASTSGSGSGLGSGSSMVSDSVSISPRDSTVTNGDDKQSSKNAVQGKRELFDGNASVRAKLAFDVALQLSNSDGKPEPKSSEALENMFNAIRPFAKLVVSDVLELDIQIENLERQNSFRPVFLNYFKTTLYHRVRAVKFRIVLAENGFDLNTLSAIWAEKNIEGLDIALQSIGRLKSKDKAELLELLDSYFDPKKTTVKPVVKGNSNNNNNYRRRNRRGGRQSVKDARPSSSPSASTEFGAGGDKSRSVSSLPDSTTKTPSPNKPRMHRKRNSRQSLGATPNGLKVAAEISSSGVSELNNSAPPAVTISPVVAQPSPTPVAITASN</u>*

### Expression and purification of Exu protein constructs

#### a) GFP-His_6_::sExu_531_

The pHTP9 plasmid vector containing the GFP-His6::sExu_531_ DNA was transformed into *E*.*coli Rosetta* cells. The protein was produced by IPTG induction at 25ºC. After an overnight (∼16h) incubation, the cells were harvested and resuspended in lysis buffer (50mM K-HEPES pH 7.5, 500 mM NaCl, 0.2% Tween 20, 30 mM imidazole, supplemented with 1 mM PMSF, protease inhibitors (Roche) and 10U of DNAse type I (NZYTech). The cells were lysed using a Emulsiflex C5 High Pressure Homogenizer at 1500 psi and clarified by centrifugation at 18000 rpm for 60 minutes at 4ºC (SorvallRC5, SS34). For purification of the fusion protein, the supernatant was loaded in an ÄKTA Pure purification system (Cytiva) onto a 5 ml HiTrap Chelating HP (GE Healthcare) charged with 0.1 mM NiCl_2_ and equilibrated with wash buffer (50 mM K-HEPES pH7.5, 500 mM NaCl, 30 mM imidazole). Extensively washed with this buffer and eluted with elution buffer (50 mM K-HEPES pH7.5, 500 mM NaCl, 500 mM imidazole), throughout a gradient of 8 CV. The fractions collected at the elution peak were analysed in a 10% acrylamide SDS PAGE at 120V for 60 min. Selected fractions were then pooled together and buffer exchanged using a PD10 desalting column (GE Healthcare) into IEX Equilibration buffer (50 mM HEPES pH7.5, 100 mM NaCl, 1 mM TCEP, 1 mM EDTA, 1 mM MgCl_2_) and loaded onto a MonoS 5/50 GL (GE Healthcare). Elution done with 50mM HEPES pH 7.5, 1M NaCl, 1mM TCEP, 1mM EDTA, 1mM MgCl_2_ in a 2-step linear gradient (10 CV up to 50%, 5CV at 50%, 10CV up to 100%). Collected fractions analysed by a 10% acrylamide SDS PAGE. Selected fraction with pure fusion protein was concentrated and buffer exchanged using an Amicon® Ultra Centrifugal Filter Unit 50MWCO (Millipore) into Extract Compatible Buffer (10 mM HEPES pH7.8, 100 mM KCl, 1 mM MgCl_2_) supplemented with 10% glycerol. Estimated concentration with NanoDrop2000 UV-Vis spectrophotometer (ThermoFisher) at 4 mg/ml, aliquoted and stored at -20ºC.

#### b) His_6_-mCherry::sExu_531_

The pET28a plasmid vector containing the His6-mCherry::sExu_531_ DNA was transformed into *E*.*coli BL21* cells for protein overexpression. The protein was produced by IPTG induction at 25ºC. After 4h incubation, the cells were harvested and resuspended in lysis buffer (50mM K-HEPES pH 7.8, 500 mM NaCl, 0.2% Tween 20, supplemented protease inhibitors (Roche) and 20U of DNAse type I (NZYTech). The cells were lysed using a Emulsiflex C5 High Pressure Homogenizer at 1500 psi and clarified by centrifugation at 18000 rpm for 60 minutes at 4ºC (SorvallRC5, SS34). For purification of the fusion protein, the supernatant was loaded in an ÄKTA Pure purification system (Cytiva) onto a 5 ml HiTrap Chelating HP (GE Healthcare) charged with 0.1 mM NiCl_2_, equilibrated with wash buffer (50 mM K-HEPES pH7.8, 500 mM NaCl, 30 mM imidazole), extensively washed with this buffer and eluted with elution buffer (50 mM K-HEPES pH7.8, 500 mM NaCl, 500 mM imidazole), throughout a gradient of 12 CV. The fractions collected at the elution peak were pooled together and loaded into a MonoS 5/50 GL column (GE Healthcare) previously equilibrated with IEX Equilibration buffer (50 mM HEPES pH7.8, 100 mM NaCl, 1 mM TCEP, 1 mM EDTA, 1 mM MgCl_2_) and extensively washed (72 CV). Elution done with 50mM HEPES pH 7.8, 1M NaCl, 1mM TCEP, 1mM EDTA, 1mM MgCl_2_ in a 2-step linear gradient (17 CV up to 50%, 15CV at 50%, 10CV up to 100%). Collected fractions analysed by a 10% acrylamide SDS PAGE. Selected fraction were pooled together to dialyse overnight at 4ºC against Extract Compatible Buffer (10 mM HEPES pH7.8, 100 mM KCl, 1 mM MgCl_2_) supplemented with 10% glycerol. Fusion protein solution was concentrated using a pre-soaked Amicon® Ultra Centrifugal Filter Unit 50MWCO (Millipore) and concentration estimated with NanoDrop2000 UV-Vis spectrophotometer (ThermoFisher) at 3.5 mg/ml, aliquoted and stored at -20ºC.

All purifications were performed using the ÄKTApure purification system (Cytiva) and the chromatographic profile of GFP::sExu_531_ protein was followed by measuring the absorbance at 280 nm, 254 nm and 488 nm, and for mCherry::sExu_531_ protein we followed by measuring the absorbance at 280 nm, 254 nm and 547 nm, in the UV-900 monitor.

### Size exclusion chromatography (SEC)

Using a ÄKTApure purification system (Cytiva), 500 µl of protein solution in Extract Compatible Buffer (10 mM K-HEPES pH 7.8, 100 mM KCl, 1 mM MgCl2 + 10% glycerol) was injected manually into a Superdex 200 Increase 10/300 GL (GE Healthcare) matrix column at a constant flowrate of 0.4ml/min. Eluted fractions of 500 µl were collected and analysed on a 10% acrylamide SDS PAGE.

The size exclusion of the column was calibrated using protein standards (Bio-Rad, catalogue # 1511901). The column had a total volume *V*_*T*_ = 24 ml and a ‘dead’ volume *V*_*0*_ = 7.2 ml. Using regression analysis, we obtained a calibration curve as follows:

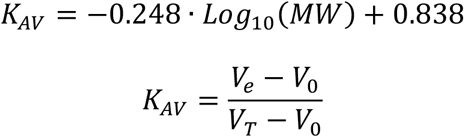

where *V*_*e*_ is the elution volume in the chromatogram. With this calibration, we expect GFP::sExu531 elution at ∼13.2 ml as monomer, ∼12 ml as a dimer and ∼11.2 as a trimer.

### Fly strains

w^1118^; hsp83-MCP::mCherry; + (gift from A. Ephrussi, EMBL Heidelberg,)

w^1118^ (Bloomington Stock Center)

w^1118^; +; p[Cas,NGE]3 (gift from T. Hazelrigg, Columbia University)

### Generation of bcdMS2(24x) transgenic fly line

Transgenic flies were generated by standard transposon-mediated transformation (Rubin and Spradling, 1982; Häcker et al., 2003). The gene of interest was first subcloned between P-element ends (pCaSper bicoid-MS2(24x)). The transgene-containing DNA was then injected into the posterior cytoplasm of young embryos (aprox. 300 embryos) in the presence of P-element transposase activity. It is expected that the DNA enters pole cell nuclei, which give rise to the future germline, where it is integrated into the genome through the action of the P-transposase. After eclosion, the injected flies (F0) were crossed and their progeny (F1) was selected for positives by means of an eye color marker present on the pCaSper vector. The insertion was mapped on the third chromosome. Positive F1 males were crossed with virgins of a 3^rd^ chromosomal balancer fly strain.

### Cleaning of coverslips

22x22 mm No. 1.5 coverslip (Marienfeld) were placed in a ceramic rack, placed into a beaker with NaOH (3M) and sonicated for 10 min. The rack was dipped-and-drained in a beaker with MilliQ water, transferred to clean MilliQ water and sonicated for 10 min. Finally, the rack was transferred into a new beaker with clean MilliQ water and sonicated for another 10 min. Coverslips were spin-dried and stored in a clean rack and sealed container until final use.

### Experimental sample preparation

Ovaries were dissected from females in a drop of halocarbon oil (Voltalef 10S, Arkema) placed on a clean coverslip, using tweezers to separate individual germaria. For some preparation that required longer time-lapse imaging, ovaries were dissected in a drop of Schneider’s medium supplemented with 10% FBS and 200 μg/mL insulin. Dissected ovaries were incubated 2x30s in 20 μL of supplemented Schneider’s medium. Finally, ovaries were transferred to a drop of supplemented Schneider’s medium on a clean coverslip next to a drop of halocarbon oil and individual germaria were pulled into the oil. This latter protocol improved the sample lifetime and allowed for longer time-lapse imaging.

### Micropipette production

Micropipettes were generated by pulling glass filaments (ID: 0.5mm, OD: 1mm, Sutter Instruments) on a vertical pipette puller (Narishige PC-10), using a one-step pulling protocol and 70% heating power. The needle was forged on a glass microforge (Narishige MF-900).

### Micromanipulation and single-cell injection

Microinjections were performed on the confocal microscope that enabled high-resolution imaging. An inverted Nikon Ti-E stage hosted four motorized micromanipulators (MPC-200, Sutter Instruments) with glass pipette holders attached to them. Motorized micromanipulation has the advantage that fast manoeuvres towards pre-defined positions can be performed with little human manual intervention. Micropipettes were back-filled with reagent, mounted on the pipette holder, and the pipette tip was positioned near the lateral side of the egg chamber. After optimization, we found that in order to reach the cytosol of nurse cells in the egg chamber the micromanipulation procedure needed to be performed with the following two steps: 1) perforation of the follicular tissue by slight lateral back-and-forth movement of the pipette tip, which promoted tisuue perforation, 2) penetration through the nurse cell membrane in direction of the needle axis, performed by fast forward movement. We note that this tissue is highly elastic and perforation is only possible if the sample is slightly immobilized. We distinguished L1 nurse cells which are in contact with the oocyte, from L2 or L3 which are anterior of L1 cells and only contact other nurse cells. Delivery of reagents into the cytosol was performed by pulse-injection using a foot pedal–controlled microinjector (FemtoJet, Eppendorf). Whenever protein samples for injection are diluted to a target concentration we used the same embryo explant compatible buffer for dilution (see above).

### Microscopy

Imaging was performed on a Nikon Eclipse Ti-E double-stage microscope equipped with a Yokogawa CSU-W Spinning Disk confocal scanner and a piezoelectric z-stage (737.2SL, Physik Instrumente), using a 20x 0.75NA multi-immersion, a 40x 1.15NA water immersion or a 60x 1.2NA water immersion objective. A laser combiner with 405nm, 488 nm and 561 nm laser lines enabled excitation of blue, green, or red fluorescence, respectively. An Andor iXon3 888 EMCCD camera with a 2x post-magnification lens was used for time-lapse acquisitions, which were programmed by Andor IQ3 software (Andor Technologies). Experiments were performed at 25ºC room and microscope stage temperature.

### Image analysis

Microscope images shown in the figures were made in Fiji/ImageJ (Schindelin et al., 2012) by calculating the maximum z-projection or the median z-projection where signal distribution was highly skewed.

## Acknowledgements

We thank the staff of the Insect Facility, the Advanced Imaging Facility (AIF) and the Technical Support Service at the Instituto Gulbenkian de Ciência (IGC). We thank L.G. Guilgur for help making a transgenic fly. Transgenic fly stocks were obtained from D. Hazzelrigg (Columbia University, NY), A. Ephrussi (EMBL Heidelberg), and the Bloomington Drosophila Stock Center (Bloomington, IN). We acknowledge M. Pinho (ITQB Nova University, Oeiras Portugal) for providing vector plasmids. This work was financially supported by Fundação para Ciência e a Tecnologia (FCT) project grant PTDC-BIA-BQM-31843-2017. We acknowledge FCT grants IF/00082/2013 supporting I.A.T.; LISBOA-01-0145-FEDER-007654 supporting IGC’s core operation; LISBOA-01-7460145-FEDER-022170 (Congento) supporting the Fly Facility; PPBI-POCI-01-0145-FEDER-022122 supporting the AIF, all co-financed by FCT and Lisboa Regional Operational Program (Lisboa2020) under the PORTUGAL2020 Partnership Agreement (European Regional Development Fund).

## Author contributions

D.V.V. and I.A.T. conceived the study. D.V.V and J.C. designed and optimized the micro-manipulation and injection experiments; R.C and D.V.V generated the mCherry::Exu construct. D.V.V. carried out all other cloning and purification experiments and biochemical analysis. D.V.V and J.C. performed microinjections. I.A.T. designed the quantification of imaging data; I.A.T. designed and assembled the microscopy setup; D.V.V and I.A.T. wrote the manuscript. D.V.V and I.A.T. jointly supervised the project.

